# Unconstrained coevolution of bacterial size and the latent period of plastic phage

**DOI:** 10.1101/2021.12.19.473366

**Authors:** Juan A. Bonachela, Melinda Choua, Michael R. Heath

**Affiliations:** Department of Ecology, Evolution, and Natural Resources, Rutgers University, New Brunswick, NJ 08901, USA; Blue Remediation Ltd., Glasgow, Scotland, UK; Marine Population Modelling Group, Department of Mathematics and Statistics, University of Strathclyde, Glasgow, Scotland, UK

**Keywords:** bacteriophage, viral plasticity, eco-evolutionary dynamics, coevolution, host-virus model

## Abstract

Viruses play critical roles in the dynamics of microbial communities. Lytic viruses, for example, kill significant proportions of autotrophic and heterotrophic microbes. The dynamic interplay between viruses and microbes results from an overlap of physiological, ecological, and evolutionary responses: environmental changes trigger host physiological changes, affecting the ecological interactions of host and virus and, ultimately, the evolutionary pressures influencing the two populations. Recent theoretical work studied how the dependence of viral traits on host physiology (viral plasticity) affects the evolutionarily stable host cell size and viral infection time emerging from coevolution. Here, we broaden the scope of the framework to consider any coevolutionary outcome, including potential evolutionary collapses of the system. We used the case study of *Escherichia coli* and T-like viruses under chemostat conditions, but the framework can be adapted to any microbe-virus system. Oligotrophic conditions led to smaller, lower-quality but more abundant hosts, and infections that were longer but produced a reduced viral offspring. Conversely, eutrophic conditions resulted in fewer but larger higher-quality hosts, and shorter but more productive infections. The virus influenced host evolution decreasing host radius more noticeably for low than for high dilution rates, and for high than for low nutrient input concentration. For low dilution rates, the emergent infection time minimized host need/use, but higher dilution led to an opportunistic strategy that shortened the duration of infections. System collapses driven by evolution resulted from host failure to adapt quickly enough to the evolving virus. Our results contribute to understanding the eco-evolutionary dynamics of microbes and virus, and to improving the predictability of current models for host-virus interactions. The large quantitative and qualitative differences observed with respect to a classic description (in which viral traits are assumed to be constant) highlights the importance of including viral plasticity in theories describing short- and long-term host-virus dynamics.

## Introduction

Viruses are the most abundant biological entities on Earth, yet cannot replicate without the synthesis machinery of a cellular host [1]. This dependence between viral replication and host is evident in the description of the infection cycle of lytic phages [2]. The lytic cycle starts when freely diffusing viruses encounter a host and attach to receptors at the cell surface. The virus then injects its genetic material into the host and hijacks the synthesis machinery to produce the components of what will be the new virions. Virions are then assembled and, finally, the expression of the holin gene leads to the perforation of the cell membrane and cell lysis, releasing the viral offspring into the medium. Key reproduction-limiting steps in the lytic cycle are the adsorption rate of viruses onto host receptors, duration of the synthesis (or eclipse) period and the virion assembly period, and the burst size (or offspring number). The time between adsorption and lysis is referred to as the latent period. Although it is still unclear what triggers lysis, all things being equal longer infection times should result in larger viral production [3].

The intertwinement between every single stage of the viral lytic infection and host metabolism and machinery [4] means that the host physiological state necessarily influences the value of these important viral traits, and therefore viral performance [5, 6, 7, 8, 9, 10]. This dependence of viral traits on host physiology (viral plasticity hereon), observed across systems, has been quantified systematically for the bacterium *Escherichia coli* and T viruses using host growth rate as a measure of physiological state [5, 6, 7, 8]. Moreover, the qualitative shape of the dependence of viral traits on host growth rate seems to be conserved across different strains of E. coli and T viruses, as well as different experimental conditions [11], thus suggesting that these functional forms stem from fundamental mechanisms generically present across systems.

In spite of the importance of viral plasticity, only recently have theories started to take it into account when exploring host-virus dynamics from an ecological [11, 12, 13] or an evolutionary perspective [11, 14, 15].

From an ecological point of view, considering viral plasticity leads to the counter-intuitive result that lower nutrient conditions are somewhat beneficial for the host, as the consequent deterioration in physiological state is compensated by the associated decrease in viral performance and reduced viral top-down pressure [11]. From an evolutionary point of view, we recently explored how viral plasticity affects the coevolution of host size and viral latent period [14]. Our results, restricted to evolutionarily stable states in which both host and virus coexist, showed that such coevolution leads to a negative correlation between host size and viral latent period. Poor growth conditions for the host selected for small radii and long infections, whereas good growth conditions selected for large hosts and short infections. For all growth conditions, host size resulting from coevolution was larger than in the absence of the virus, and the viral latent period was longer than if hosts did not evolve. Here, we revisit the system to expand our theoretical framework and explore a wider range of potential evolutionary outcomes, including the possibility for evolutionary branching and the possibility for coevolution to drive either host or virus to extinction (evolutionary collapses).

We aim to understand the mechanisms underlying the different outcomes of the host-virus dynamics represented in our model, and the role played by viral plasticity. These dynamics are highly nontrivial, as short generation times and vast populations of both host and virus lead to rapid evolution that, ultimately, may trigger interactions between ecological and evolutionary timescales (eco-evolutionary dynamics) [16, 17]. Thus, we will pay especial attention to the coevolutionary transient, i.e. the evolutionary path leading to such outcomes. We also extend our framework to consider a larger variance for the evolutionary step, which ensures the robustness of the evolutionary equilibrium reached by the system. In addition, we include viral avoidance of secondary infection by other viral individuals (superinfection), which not only does it increase the realism of the framework but also increases its dynamical stability by reducing the amplitude of the antagonistic demographic oscillations that characterize populations in host-virus systems. These modifications of the model aim to ensure that more replicates reach a meaningful outcome that results from the eco-evolutionary dynamics between host and virus, thus increasing the breadth and robustness of our study.

As in the previous version of the model, we will focus on cell radius as the evolving trait for the host, and latent period as the evolving trait for the virus. As justified below in detail, size is a “master trait” for microbes (e.g. [18]) that influences every aspect of their physiology and ecological interactions, thus affecting most other host traits. On the other hand, the latent period is one of the most important life-history traits for the virus, and an ideal trait to study in an evolutionary framework given the small pleiotropic effects on other traits [19].

Understanding not only stable coexistence but also branching and collapse, and how those equilibria were reached, can potentially provide key information about a variety of systems. For example, it can help us understand the eco-evolutionary dynamics governing phytoplankton blooms, which end with very low phytoplankton densities or even collapse, an outcome in which viruses play a prominent role [20, 21, 22]. Including plasticity can improve the dynamic description of other viral-mediated processes that are key for biogeochemical cycles and microbial diversity (e.g. viral shunt and shuttle) [22, 23, 24], and therefore the reliability of models that estimate primary production. The dynamics, and not only the end result, may also be of relevance to design anti-bacterial treatments (phage therapy [25]), or when studying viral infections of changeable communities such as the animal microbiome. Finally, including viral plasticity can be key to reliably representing the viral infection of biofilms, which are characterized by a very variable (temporal and spatial) distribution of nutrient availability, and thus by a mosaic of growth conditions within and around the biofilm [26, 27].

## Methods

To explore the unconstrained coevolutionary dynamics of host and virus, we used a standard delay model that has been shown to produce realistic ecological and evolutionary behavior [11, 28, 29]. This model describes the dynamics of the population of uninfected hosts (*C*, in *cell* · *l*^−1^), infected hosts (*I*, in *cell* · *l*^−1^), free viruses (*V*, in *ind* · *l*^−1^), and the concentration of the most limiting nutrient for the host (*N*, in *mol* · *l*^−1^), all interacting in a chemostat environment:

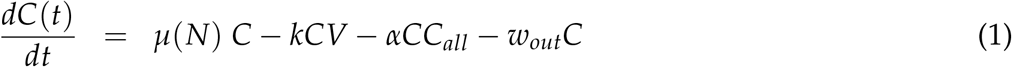

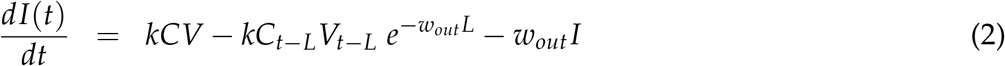

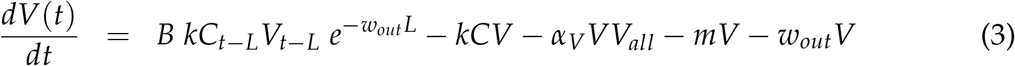

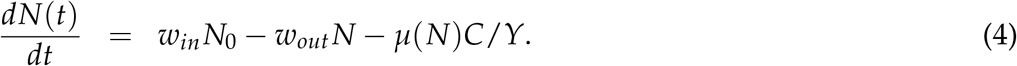

See table 1 for symbols and units. The population of uninfected hosts increases thanks to the uptake of the most limiting nutrient (here, glucose; first term, Eq.(1)), declines due to infection (second term) or dilution from the chemostat (last term). Similarly to [14], we consider here potential competition for space, light, or other resources not explicitly modeled that can affect negatively the growth of the focal population (third term), a plausible scenario as the framework introduces in ecological time new host mutants (see below); here, *C*_*all*_ represents all hosts from all phenotypes in the system.

**Table 1:** Symbols for variables used in the model and parameter values. Data for the host obtained from [55, 56, 57], and host tradeoff functions and their parametrizations from [33, 36, 37]; data for the virus into the ranges shown/used in [58, 59, 60, 44]. For the calculation of the yield factor from the references, we assumed a fixed carbon content per host cell of 10^−12^ *g*. For other conversions, we used that the carbon content of glucose is 180.15588 *mol* · *mol*^−1^.

| Symbol | Description | Units | Value |
| --- | --- | --- | --- |
| $L$ | Latent period | $d$ | Evolutionary variable |
| $r$ | Host cell radius | $\mu\text{meter}$ | Evolutionary variable |
| $C$ | Density noninfected host | $\text{cell} \cdot l^{-1}$ | Eq.(1) |
| $I$ | Density infected host | $\text{cell} \cdot l^{-1}$ | Eq.(2) |
| $V$ | Density free virus | $\text{ind.} \cdot l^{-1}$ | Eq.(3) |
| $N$ | Glucose concentration | $\text{mol} \cdot l^{-1}$ | Eq.(4) |
| $\mu$ | Host growth rate | $d^{-1}$ | Eq.(5) |
| $\mu_{max}$ | Maximum growth rate for host | $d^{-1}$ | Eq.(6) |
| $K_N$ | Half-saturation constant for growth | $\text{mol} \cdot l^{-1}$ | Eq.(8) |
| $k$ | Adsorption rate | $l \cdot \text{cell}^{-1} \cdot d^{-1}$ | Eq.(7) |
| $B$ | Burst size | *virions* | Eq.(9) |
| $E$ | Eclipse period | $d$ | Eq.(10) |
| $M$ | Maturation rate | *virions* $d^{-1}$ | Eq.(11) |
| $\mu_{max_{exp}}$ | Max. host growth rate from experiments | $d^{-1}$ | 40.8 |
| $\mu_{max_{ref}}$ | Saturating max. growth rate in Eq.(8) | $d^{-1}$ | 28.8 |
| $K_{N_{ref}}$ | Half-saturation constant at $\mu_{max} = 0$ | $\text{mol} \cdot l^{-1}$ | $3.55 \cdot 10^{-8}$ |
| $D_0$ | Diffusivity T-like viruses in water | $\text{dm}^2 \cdot d^{-1}$ | $3.73 \cdot 10^{-5}$ |
| $Y$ | Yield parameter | $\text{cell} \text{ mol}^{-1}$ | $9 \times 10^{13}$ |
| $m$ | Virus mortality rate | $d^{-1}$ | 0.09 |
| $N_0$ | Glucose input/supply concentration | $\text{mol} \cdot l^{-1}$ | $3 \cdot 10^{-6}, 5 \cdot 10^{-6}, 10^{-5}, 5 \cdot 10^{-5}, 10^{-4}$ |
| $w_{out}$ | Chemostat dilution rate | $d^{-1}$ | $0.05\mu_{max_{ref}} - 0.55\mu_{max_{ref}}$ |
| $w_{in}$ | Chemostat inflow rate | $d^{-1}$ | $w_{out}$ |
| $a$ | Factor of Eq.(6) | $d^{-1} \cdot \mu\text{m}^{-3p}$ | $(24 \cdot 10^{-12}) \cdot 10^{1.9}$ |
| $p$ | Exponent of Eq.(6) | (-) | 0.177 |
| $dr$ | Min. difference in $r$ for host phenotypes | $\mu\text{meter}$ | 0.01 |
| $dL$ | Min. difference in $L$ for viral phenotypes | $d$ | 0.003472 |
| $\sigma_r$ | Std. dev. host evolutionary step | $\mu\text{meter}$ | $2dr = 0.02$ |
| $\sigma_L$ | Std. dev. viral evolutionary step | $d$ | $2dL = 0.00694$ |

The population of infected individuals increases due to infection (first term, Eq.(2)), and declines when infected cells are diluted (third term) or lysed (second term). Note that the number of cells that are lysed at time *t* were infected a latent period *L* in the past, and thus this lysis term considers the number of infections at time *t* − *L* weighted by the probability to survive dilutions in that period 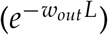. Cells lysed at time *t* produce the new batch of viruses (first term of Eq.(3)), and the population declines as viruses infect new hosts (second term), decay (become non-infective after a period of time, fourth term), or are diluted from the chemostat (last term). We expanded the model from [14] to include also the possibility for phage to avoid superinfection, thus accounting for the battery of mechanisms that a phage that has entered the host can deploy to prevent any other virus from using the same host for replication [30]; this mechanism, here implemented with a density-dependent term (third term, where *V*_*all*_ represents the sum of all viral density across phenotypes), is not only more realistic but also reduces the mathematical instability reported in [13, 14]. Finally, the dynamics of the most limiting nutrient (glucose) include the inflow and dilution of nutrient that characterize the chemostat environment (first and second terms), and the uptake of the nutrient by the uninfected hosts (last term). We assumed that cell growth and replication stop at infection, reason why infected cells do not take up glucose and why they are not considered for the density-dependent term (third term) in Eq.(1). Nutrient uptake is calculated based on requirement for growth, with the growth rate given by the classic Monod formulation [31]:

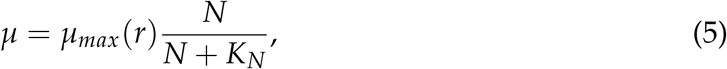

where *µ*_*max*_ is the maximum growth rate (in *d*^−1^), and *K*_*N*_ (in *mol* · *l*^−1^) the half-saturation constant for growth on nutrient *N*.

Following standard modeling practice, we set a threshold below which either the total host (i.e. infected and uninfected cells) or the free viral population are considered to be extinct. This practice prevents unrealistically low values of the population densities from “regenerating” populations. Here, we set a threshold of 1 *ind* · *l*^−1^ for either population. Differently from [14], we consider that the threshold for the virus includes not only free viruses but also replicating viruses (i.e. viruses that are currently infecting a host), which prevents the elimination of viral mutant populations that entered the system recently and are still inside infected cells as part of their first lytic cycle. Neither reverting this more conservative rule, nor eliminating the new superinfection avoidance term in Eq.(3), altered qualitatively our results.

### Traits and trade-offs

Although the model above can be applied to any bacteria-phage system, for the sake of concreteness here we consider the dynamics of one of the most studied examples: T phage infecting *Escherichia coli*. See table 1 for parameter values.

As explained in detail in the next section, we focus on size as single evolutionary trait for the host. The choice is justified because, for microbial organisms, size is linked to almost every aspect of its life cycle and ecological functioning (e.g. [18, 32]). Particularly for E. coli, it has been shown that the maximum growth rate is positively correlated with cell size [33]:

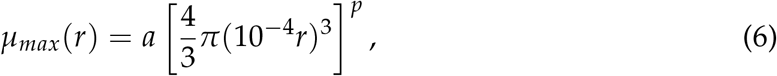

where we used the spherical approximation for the cell, *r* is cell radius (in microns), and *a* and *p* are parameters shaping this power-law correlation (see table 1 for values and units). Cell size also affects the rate of encounters with the virus, which can be accounted for using the following expression for the adsorption rate, *k* (in *l* · *d*^−1^) [34]:

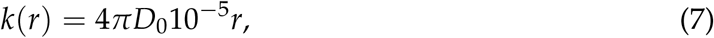

with *D*_0_ (in *dm*^2^ · *d*^−1^) the diffusivity of T-like viruses in water. Since T viruses use lipopolysaccharides (LPS) as main target receptors for adsorption to the E. coli cell, and LPS are very abundant on the cell surface (more than 75% coverage) [35], we assumed for simplicity that all encounters led to a successful adsorption.

In addition, we considered the following correlation between the half-saturation constant for growth and the maximum growth rate [36, 37]:

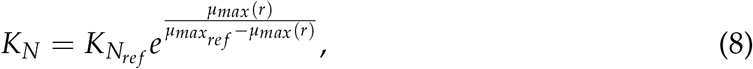

which effectively links the half-saturation constant with size as well. Thus, large cell radii lead to high growth potential but low affinity (inverse of *K*_*N*_), therefore setting a tradeoff for the host. The parameter 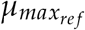 represents a maximum value for *µ*_*max*_, and 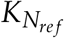 is the half-saturation value for *µ*_*max*_ = 0 (see table 1 for values).

Together with the adsorption rate *k* above, the main viral traits that define the infection cycle are the eclipse period, *E* (in *d*), the maturation rate, *M* (in *virion* · *d*^−1^), the burst size, *B* (in *virion* · *cell*^−1^), and the latent period, *L* (in *d*). Here, we considered the latter the focus of viral evolution, since it determines the timing of lysis and limits virion production by setting a maximum time for synthesis and assembly [10]. Such a limitation results in a correlation between burst size and latent period, under the assumption that the timing to exhaust intracellular resources is longer than the timing of lysis [3]:

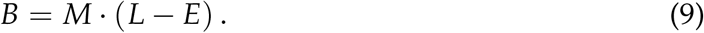

As explained above, these viral traits are affected by the host physiological state. Here, we used the host growth rate to represent the cell’s physiological state because light, temperature, nutrient availability, and other factors affecting host physiology are ultimately reflected in changes in growth rate. This variable has been used in the past to study how T viruses respond to changes in host physiology by exploring how the value of different viral traits depends on host growth rate [5, 6, 7, 8]. An effort to characterize these relationships across E. coli-T phage systems showed that the eclipse period can be expressed as [11]:

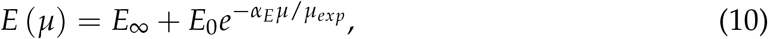

and the maturation rate as:

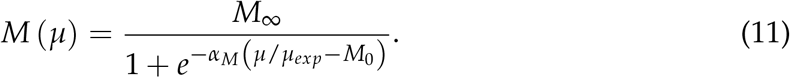

In short, the eclipse period decreases and the maturation rate increases as a function of host growth rate. The maturation rate increases as a sigmoid and reaches a plateau for high host growth rates. The eclipse period decreases exponentially from a non-zero maximum value for *µ* → 0 to a minimum value for *µ* → *µ*_*exp*_. The latter is the maximum growth rate observed in the experiments of reference [6] (see table 1). The former indicates the possibility for phage to replicate even for negligible host growth, which has been observed experimentally for this system [7]. Here, we focused on obligate lytic viruses only, but an alternative strategy for the virus in such challenging conditions is to use a temperate replication mode and switch to lytic mode when appropriate (e.g. [38]).

We refer to [11] for further details and biological justification of these functional forms.

From Eqs.(10)-(11), it follows that the burst size will be affected by the host growth rate (Eq.(9)), which has been observed experimentally in the past (e.g. [8]). The timing of lysis is also influenced by host physiological state; since *L* is the focus of viral evolution here, however, we let coevolution with the host determine its value and how it depends on the host growth rate.

In contrast, models that do not consider viral plasticity use fixed values for the viral traits above, typically obtained from experiments in which the host is grown at optimal conditions [5]. To study the effect that accounting for plasticity has on standard predictions for host-virus dynamics, in previous work we parametrized the nonplastic case by setting viral traits to their value for best growth physiological status, i.e. by using Eqs.(10)-(11) with *µ* = *µ*_*max*_ [11, 13, 14]. In [14], however, host maximum growth rate was affected by the evolution of host size; thus, although setting *µ* = *µ*_*max*_(*r*) was technically consistent with the fact that nonplastic parametrizations rely on “best host growth” values, it somewhat allowed for a form of viral plasticity because the same viral population infecting different host phenotypes (i.e. with different *r*) would show different trait values. Here, we ensured that the nonplastic case fulfills both aspects of the definition above by using Eqs.(10)-(11) with the maximum value for the host growth rate allowed by the tradeoff expression, Eq.(8). In other words, 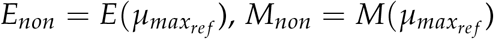 and *M*_*non*_(*L* − *E*_*non*_), which ensures fixed traits for a given viral phenotype regardless of the host it infects.

### Evolutionary dynamics

We embedded the model above in an evolutionary framework that has been successfully used in the past to study various aspects of microbial evolution (e.g. [11, 29, 39]). Differently from other traditional evolutionary frameworks, this framework does not impose a separation of the ecological and evolutionary timescales. Instead, it allows for mutations to occur at random times, and thus for new mutant phenotypes to be introduced in the system at ecological timescales.

Our focus on host size and viral latent period as only evolving traits means that these traits characterize host and viral phenotypes, respectively. Thus, all host phenotypes were identical except by their size (and related traits, Eqs.(6)-(8)); similarly, given a host growth rate, viral phenotypes only differed in the value of the latent period (and, thus, the burst size, Eq.(9)).

The system was initialized with a single host and viral phenotype using a random value for *r* and *L* respectively. These initial populations interacted through Eqs.(1)-(4). At mutation events, the mutating phenotype was selected at random following a roulette-wheel algorithm in which the phenotype with the highest population density (hereon “dominant phenotype”, for either host or virus) had the highest probability to be chosen. Thus, a population of a new phenotype was introduced whose trait values were identical to those of the parent except for the evolving (and related) traits. The mutant’s value for the evolving trait was chosen at random following a Gaussian distribution centered around the value of the parental trait, and standard deviation given by *σ*_*r*_ (for the host) or *σ*_*L*_ (for the virus). We assumed that two phenotypes were different only if the new value of the trait differed from any existing phenotype an amount *dr* (for the host) or *dL* (for the virus); thus, we set the standard deviation for evolutionary steps to be *σ*_*r*_ = 2*dr* and *σ*_*L*_ = 2*dL*. This choice allowed for an evolutionary step that was much less restrictive than the one used in [14], which enabled a more efficient and complete exploration of the trait space, yet was sufficiently small for the new phenotype to still be considered a mutation from the parental phenotype. Setting a fixed mutation time for host and virus (either comparable or with the virus mutating faster than the host), did not alter qualitatively our results but changed the number of competing phenotypes at any given time.

To understand the role of environmental factors on the coevolutionary outcome, we explored different values of both the nutrient input concentration, *N*_0_, and the dilution rate, *w*. For the latter, we used fractions of the maximum possible growth rate in the system, set by 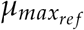 (see table 1). In addition, to understand to what extent the coevolutionary behavior of the host resulted from bottom-up versus top-down regulation, we compared the host size emerging from coevolution with results obtained in the absence of viruses. The latter provided a reference for evolutionary behavior in response to purely bottom-up processes (in this case, the availability and uptake of glucose). Moreover, we compared the latent periods emerging from coevolution with those obtained in the absence of host evolution. For the latter, we used an analytical expression for the optimal latent period obtained under the assumption that host does not evolve and viral evolution aims to minimize “host use”, as per resource competition theory [11, 40] (see Supplementary Information). This comparison allowed us to explore under which environmental conditions the viral strategy departed from “host density minimization” due to host coevolution.

We further assumed that all viruses can infect all hosts. This simplifying assumption was justified by the observation that the coevolution of bacteria and phage can lead to the emergence of generalist viruses [41]. Another potential outcome of coevolution is the possibility for the virus to lyse all available hosts after, e.g. a “host immunity” vs. “viral immunity avoidance” arms race [42], an example of evolutionary suicide through Tragedy of the Commons [43]. We investigated whether host extinction due to either an “evolutionarily underperforming” host or “evolutionarily overachieving” virus could happen through the coevolution of host size and viral latent period. Thus, in addition to the replicates that resulted in the coexistence of a clear dominant host and virus phenotype [14], we also analyzed the cases in which extinction occurred after a minimum number of days, set to 1, 000 days. Because all replicates start with one random pair of host and virus phenotypes, setting a conservative minimum period of survival for the system filters extinctions occurring due to an unstable initial condition [13] instead of due to evolutionary dynamics.

## Results

Our eco-evolutionary framework allowed for the overlapping and mutual interaction of host-virus coevolution and viral plasticity. Due to the stochastic nature of the replicates (from the initial condition to the random exploration of the trait space for both host and virus), the results below consider instances among 300 replicates that were classified according to whether a dominant host and virus phenotype coexisted at maximum time of the replicate (30, 000 days), or extinction occurred after 1, 000 days. Replicates showing the remaining option (extinction occurring before the minimum number of days) were rejected as the result of a pathological random combination of initial host and virus phenotypes.

### Host-virus coexistence

The number of replicates that showed coexistence increased with the input concentration (*N*_0_) for the plastic case, but decreased for the classic parametrization in which viral traits are fixed (“nonplastic case” hereon). Both consistently showed such surviving runs for *N*_0_ ∼ 5 · 10^−6^ − 10^−5^ *mol* · *l*^−1^. In these surviving replicates, the stochastic exploration of the trait space by both host and phage led to an evolutionarily stable strategy (ESS) given by a value for both evolving traits, host size and viral latent period. Figure 1 shows an example of how the distribution of abundances for host (left panel) and phage (right panel) phenotypes changed over time. Both host and virus show a distribution (i.e. non-negligible variance in trait value) with a well-defined dominant phenotype, and alternation of dominance over time until reaching a stationary state. Such a stationary state may show demographic oscillations, especially for high *N*_0_ or the nonplastic case (not shown). Thus, a well-defined evolutionary stationary state was reached for all surviving replicates in spite of the standard deviation for evolutionary steps, *σ*_*r*_ and *σ*_*L*_, being set to twice the minimum trait difference characterizing phenotypes (*dr* and *dL*, see Methods and table 1), emphasizing the stability of the evolutionary steady state. Figure 1 shows that the host size reached its evolutionary stationary value by following more abrupt changes and fewer alternations of dominance than the viral latent period; in addition, the host trait reached its stationary value before the viral trait, which seemed to be generally the case. The overwhelming dominance of the most abundant phenotype at the stationary state facilitated the selection of the dominant as representative of that replicate (in Fig. 1, (*r, L*) = (0.98, 0.09)).

**Figure 1:**
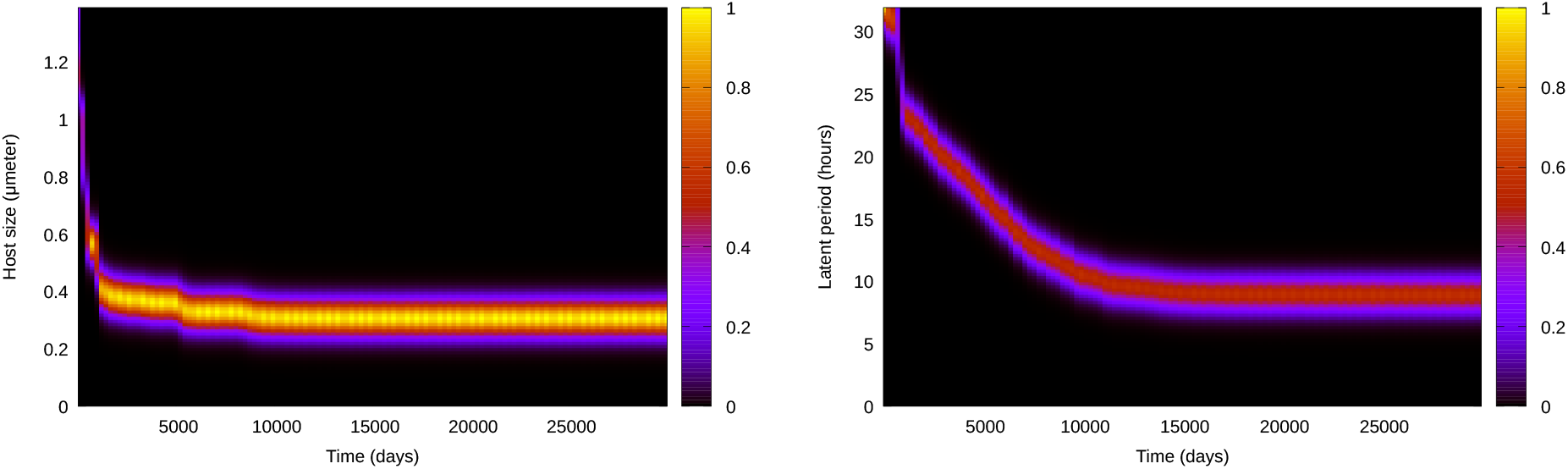
Density plot obtained with the abundance distribution at each time for all host (left) and virus (right) phenotypes, with color representing normalized density. Each population is composed of a clear dominant phenotype (value of host radius and viral latent period, respectively), and others with similar trait values. Both populations reach a stationary state that enables the definition of an evolutionarily stable strategy (ESS) for both evolving traits. Simulations obtained for the plastic case with 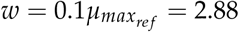 and *N*_0_ = 10^−5^ *mol* · *l*^−1^.

Due to the stochastic character of the eco-evolutionary dynamics, the stationary trait values reached across replicates were not identical. Nonetheless, a representation of the abundance of all phenotypes collected at the end of each replicate (heatmap in Fig. 2) showed that the final host and viral populations were mostly composed of a single dominant with a trait value that was similar across replicates. Thus, we did not observe evolutionary branching. In addition, we independently calculated the trait value representing the ESS for the particular environmental conditions (i.e. for a given *N*_0_ and *w*). To this end, we selected each replicate’s final dominant host and virus by calculating, for each phenotype, the median of their abundance in the last 100 days of the simulation, and identifying the phenotype with the highest median. We then calculated the ESS for the given *N*_0_ and *w* by averaging across replicates the trait value of the dominant. As the points in Fig. 2 show, these average (*r*_*ESS*_, *L*_*ESS*_) matched the most abundant trait combinations observed across replicates.

**Figure 2:**
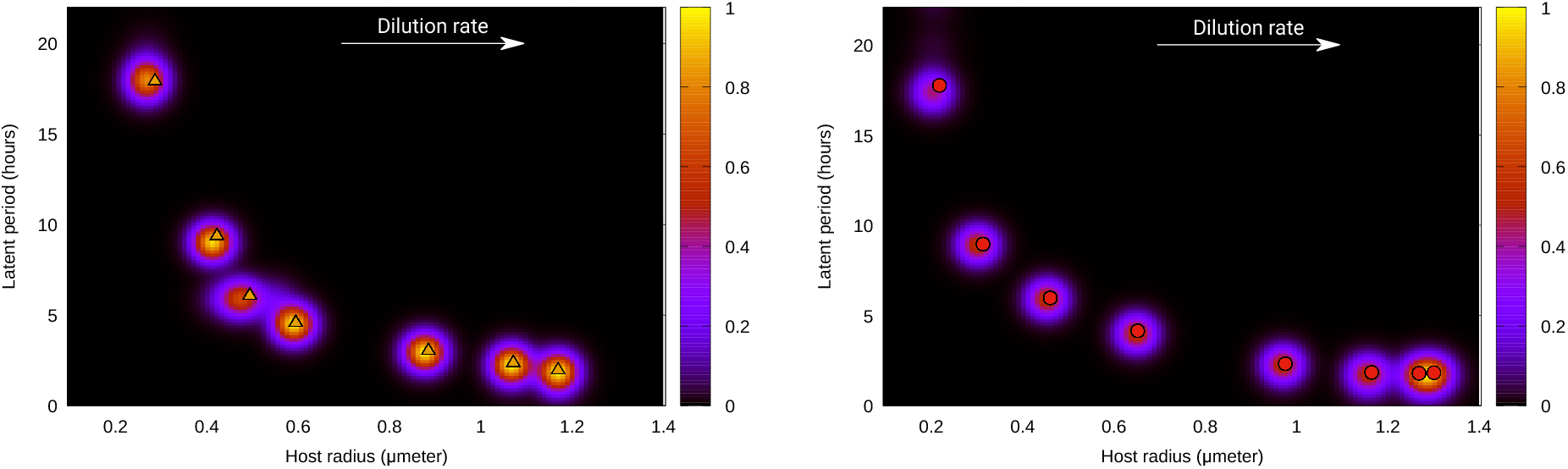
(Normalized) density plots for the final state of the system (heatmap) and average value for traits, (*r*_*ESS*_, *L*_*ESS*_) (points), for different dilution rates, *w*. Left panel, for nutrient input concentration *N*_0_ = 5 10^−6^ *mol · l*^−1^; right panel, for *N*_0_ = 10^−5^ *mol · l*^−1^. Both panels, obtained for the plastic case, show the dominance of one well-defined value for host and viral traits.

The emergent host size, *r*_*ESS*_, was positively correlated with the dilution rate, opposite to the emergent latent period, *L*_*ESS*_ (Fig. S1). For the former, a higher input concentration, *N*_0_, led to noticeably higher radii only for high *w*; the external nutrient input also somewhat increased the slope of this correlation, and thus for low dilution rates the host radius emerging under low *N*_0_ was slightly larger than under high *N*_0_. The input concentration decreased the emergent latent period. In the nonplastic case, hosts showed a smaller size and viruses showed a smaller latent period than the plastic case, especially for low *w*. Thus, for both plastic and nonplastic cases, *r*_*ESS*_ was inversely correlated with *L*_*ESS*_, and such *r*_*ESS*_ vs. *L*_*ESS*_ curve shows a down-left shift for the nonplastic case (Fig. 3, left). *N*_0_ barely affected this relationship qualitatively nor quantitatively.

**Figure 3:**
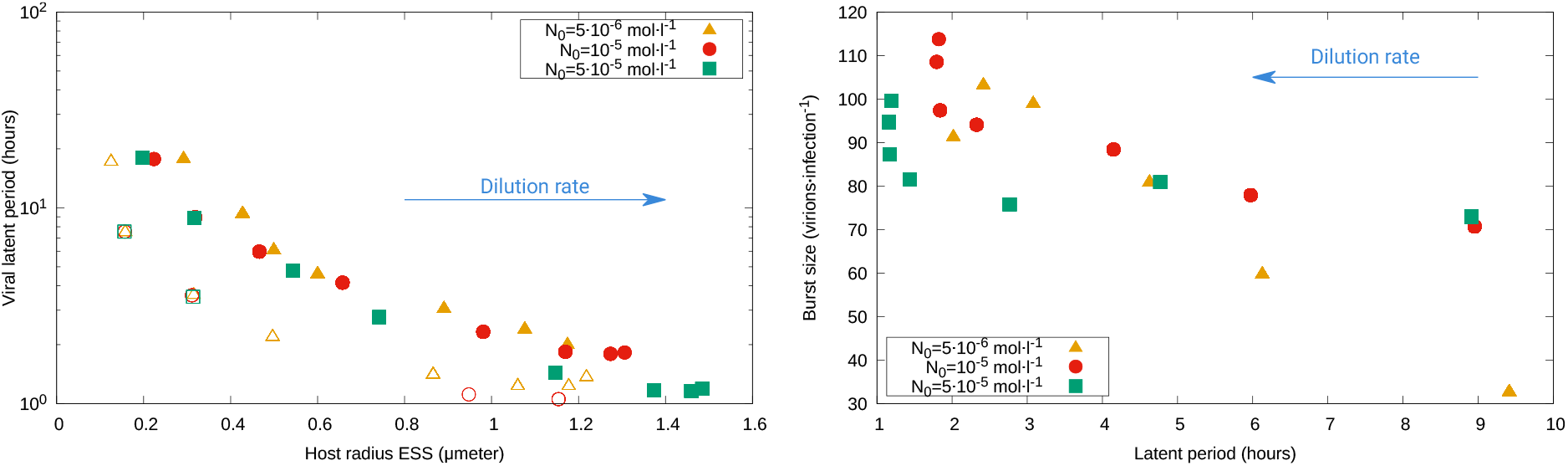
Average across replicates for different traits and cases. Left: Dominant host radius versus dominant latent period; both plastic (full symbols) and nonplastic (open symbols) cases show a negative correlation between *r*_*ESS*_ and *L*_*ESS*_, but smaller hosts and shorted infections dominate in the nonplastic case. See Fig. S1 for trait dependence with dilution rate, *w*, and Fig. S5 for other versions of the model. Right: Dominant burst size versus dominant latent period, showing a negative correlation between *B*_*ESS*_ and *L*_*ESS*_ for the plastic case that becomes steeper as *N*_0_ increases. See Fig. S3 for nonplastic case and dependence of *B*_*ESS*_ on *r*_*ESS*_.

As explained in Methods, viruses in the nonplastic case are characterized by constant eclipse period and maturation rate (Fig. S2), which leads to a constant burst size as well (Eq.(9)). As a consequence, the relationship between the viral latent period and burst size showed an opposite correlation for the plastic and nonplastic cases (Fig. 3, right panel, and Fig. S3, left). The latter conserves the classic positive correlation (and therefore tradeoff) between offspring number and infection time, whereas plasticity allows lower latent periods to reach larger burst sizes, dissimilarity reported in previous work that considered viral evolution only [11].

The presence or absence of the *B*-*L* tradeoff leads to an opposite interdependence of emergent burst size and host radius (Fig. S3, right): while the nonplastic case shows a negative correlation, the plastic case leads to a subtle positive correlation (non-monotonic for lower *N*_0_). For the nonplastic case, this decline results from a decreasing assembly period (difference between the emerging *L* and the fixed *E* value, see Fig. S4 left); for the plastic case, the decreasing assembly period (in this case defined as the difference between the emerging *L* and the *E* value set by the dominant host) is compensated by the increasing maturation rate as *w* increases.

Removing the “superinfection avoidance” term in Eq.(3) and setting a viral extinction threshold that ignores intracellular viruses and focuses on free viruses only (see Methods) did not alter qualitatively the results of the model with plasticity (Fig. S5).

The average nutrient concentration at stationarity was only noticeably impacted by the host only when both *w* and *N*_0_ were low (Fig. S4, right), with hosts drawing down nutrient to lower levels in the plastic case. The density of the dominant host showed a nonmonotonic dependence on *w*, reaching a minimum for intermediate dilution rates (Fig. 4, left). Differences in host density across nutrient input concentrations were barely noticeable for low *w*, small for large *w*, but large for intermediate *w* (e.g. more than an order of magnitude between the *N*_0_ = 5 · 10^−6^*mol* · *l*^−1^ and *N*_0_ = 5 · 10^−5^*mol* · *l*^−1^ cases). The average density of the dominant virus showed an overall decreasing trend with both dilution rate for high input concentration, and for the nonplastic case (Fig. 4, right). In the plastic case, lower *N*_0_ led to a more complicated pattern, with nonmonotonic changes that remained within the range 10^8^ - 5 · 10^9^ *cell* · *l*^−1^ for all *w*.

**Figure 4:**
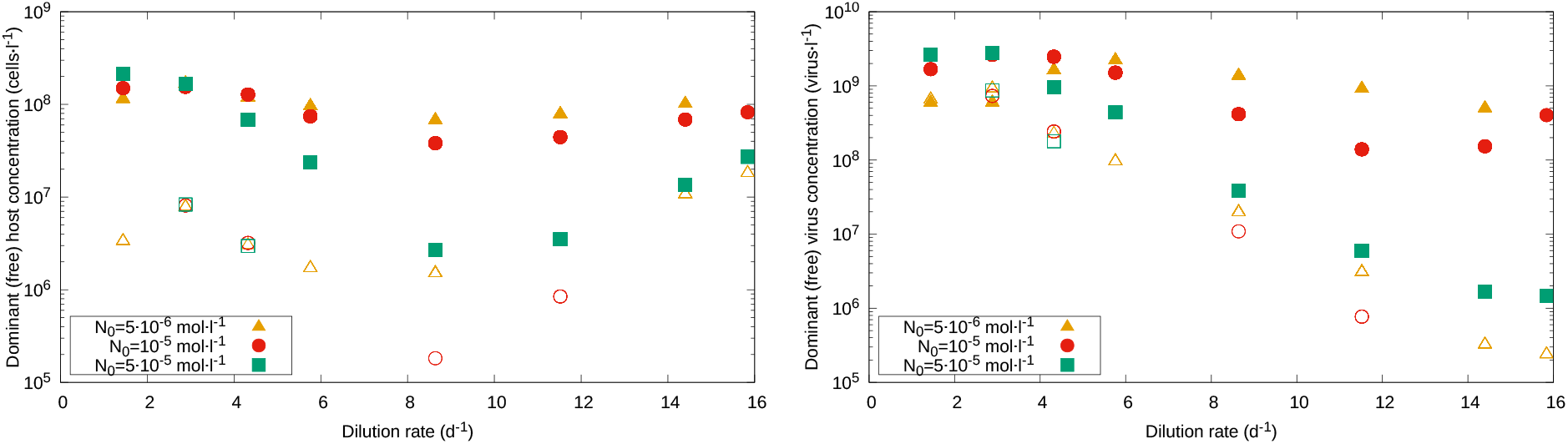
Density of the dominant population as a function of the dilution rate, *w*. Left: The concentration of the dominant host, averaged across replicates, initially decreased then increased, with a minimum value observed for intermediate *w*; concentrations were smaller for higher *N*_0_ and the nonplastic case. Right: The concentration of the dominant viral phenotype decreased with *w* for high nutrient input and the nonplastic case, but remained within the 10^8^ -5 10^9^ *cell l*^−1^ range for lower *N*_0_. Full symbols represent the plastic case, and open symbols the nonplastic case.

### Single evolution

The radii emerging in the presence of the virus were positively correlated with those emerging in the absence of it, and the range of dilution rates for which departures from the 1:1 line (i.e. differences between the *r*_*ESS*_ with or without the virus) occur depended on the input concentration (Fig. 5, left). While, for low *N*_0_, departures were constrained to low dilution rates, for high *N*_0_ differences occurred for any *w*. In all cases in which there were differences, hosts in the presence of the virus were smaller than in the absence of it.

**Figure 5:**
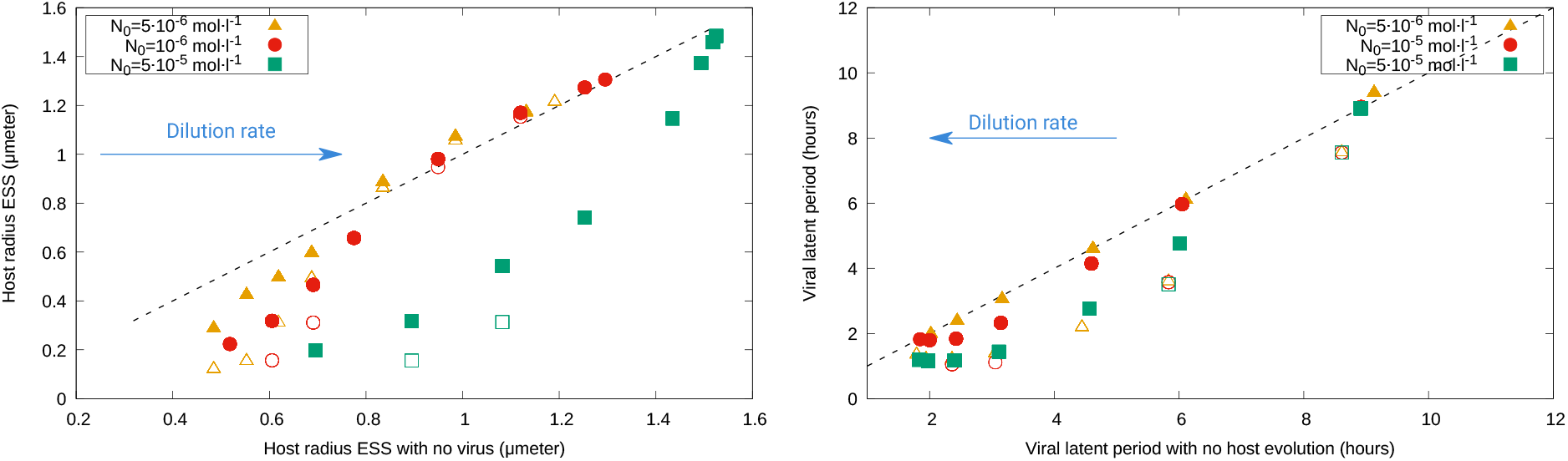
Left: Comparison between the host radius emerging from evolution with and without the virus; discrepancies (departures from the 1:1 line) occur for low dilution rates and for high nutrient input concentrations. Right: Comparison between the latent period emerging from coevolution with the host versus minimization of a fixed-radius host; differences occur for high *w*, and are accentuated by *N*_0_ in the plastic case only. As before, full symbols represent the plastic case, and open symbols the nonplastic case.

For the latent period, high dilution rates led to shorter infections than in the absence of host evolution (the latter calculated as *L*_*nocoev*_ = 1/*w*_*out*_ + *E*(*µ*), see Supplementary Information and [11, 29, 44]), but differences decreased as *w* or *N*_0_ decreased (Fig. 5, right). For any input concentration, the nonplastic case curve was similar to that of the high-*N*_0_ plastic case.

### Evolutionary collapse

As mentioned above, the number of replicates that resulted in extinction decreased with the input concentration for the plastic case and increased for the nonplastic case. We observed artificial (early) extinctions (see Methods) for almost all combinations of dilution rates and nutrient input concentrations, and only in a reduced number of cases extinctions occurred after the minimum 1, 000 days we set to discern whether the collapse resulted from the evolutionary process.

In these cases of evolutionary collapse, the host went to extinction before the virus, thus leading to the eventual collapse of the whole system. In this evolutionary path to extinction, the host seemed to be approaching the ESS corresponding to the case without the virus, while the virus was still evolving after starting from an initial phenotype with a high *L* value (Fig. S6). The radii of the host phenotypes that were present immediately before extinction showed a negative correlation with the latent period of the dominant virus (Fig. S7). The correlation was less marked in the plastic case (left panel) than for the nonplastic case (right). For the latter, the negative correlation was only broken for low radii. For the former, mid-to-low radii led to evolutionary collapse for a well-defined range of *r* (0.6 − 0.7 in Fig. S7) but a much wider range of mid-to-high *L* (≈ 20*h* in the example).

## Discussion

The vast numbers and short generation time of microbial organisms, as well as the possibility of rapidly changing environments, emphasizes the importance of theories that take into account the overlapping effects of ecological and evolutionary responses to describe the dynamics of these organisms. A reliable description of the interaction between bacteria and phages thus requires accounting for coevolutionary responses as well as the effect that changes in host physiology has on the main traits characterizing viral performance (viral plasticity).

In previous work, we explored such coevolution by focusing on host size and viral latent period as evolving traits, constraining our study to cases in which both host and virus coexisted and showed an evolutionarily stable strategy (ESS) [14]. Here, we sought to eliminate any such constraints and study any outcome of this coevolutionary process, including potential evolutionary branching and extinction. One of such constraints was the use of an infinitesimal evolutionary step [45]; the larger standard deviations *σ*_*r*_ and *σ*_*L*_ used here enabled a more thorough exploration of the trait space while keeping a reasonable phenotypic link between offspring and parent. Some of the different results observed with the current version may thus result from the current framework being able to find global evolutionary attractors and elude “evolutionary traps” (local attractors with deep basins of attraction). In addition, we sought to increase the stability of the system and reduce collapses not driven by evolution but instead by the random initial condition. To this end, we included viral superinfection avoidance, commonly observed for phages [30], through a density-dependent competition-like term in Eq.(3) that reduced the amplitude of demographic oscillations. Moreover, we considered intracellular viruses when assessing whether a phenotype fell below the extinction threshold, which prevented the removal of viral phenotypes before their first latent period ends. Finally, we used here a more strict implementation of the classic (i.e. nonplastic) case that ensured that each viral phenotype’s traits were fixed at all times regardless of the host.

Figure 6 summarizes the phenomenology we observed with our framework, and the mechanisms we hypothesize below underlie those observations.

**Figure 6:**
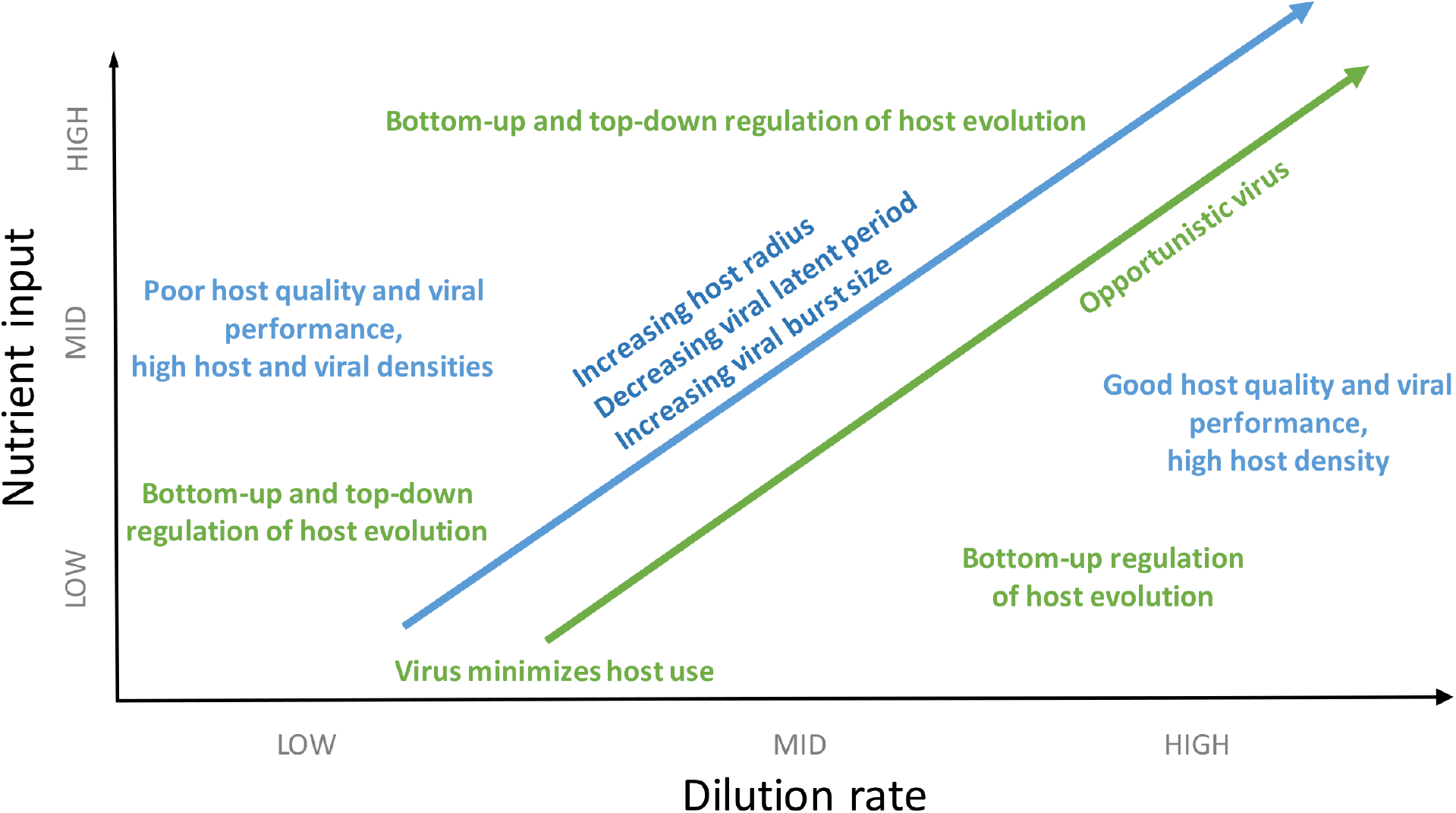
Schematic figure summarizing the phenomenology observed in the plastic case (blue tones) and hypothesized underlying mechanisms (green tones) as environmental conditions, represented by nutrient input concentration (vertical axis) and dilutionr rate (horizontal axis), change.

### Host-virus coexistence

The unconstrained evolution described by the current model led to results similar to the previous version when host and virus coexisted [14], but also to important differences.

Similarly to our previous study, the final stationary state was composed of a well-defined dominant phenotype for both host and virus. In other words, the system reached a well-defined ESS, with no instances of evolutionary branching. Nonetheless, the evolutionary path towards the ESS showed evolutionary leaps, more markedly observed for the host trait. Between leaps, distributions of the evolving traits were peaked but with a non-negligible width, with leaps being a transitory equalizing moment that enabled less-peaked distributions [46]. As expected, the evolutionary path towards the ESS in the current study showed a much wider degree of variation for both host and virus traits. Also differently, in the current model plasticity led to larger infection times than when viral traits were fixed (nonplastic case), which results from parametrizing the latter strictly using values corresponding to the best possible physiological state that any host can show (i.e. 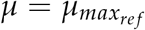) thus leading to the shortest possible latent periods.

As in [14], the emergent host size was inversely correlated with the emergent viral latent period, that is, smaller hosts selected for longer infections. Moreover, the potential evolutionary payoffs for these strategies also depended on environmental conditions. For example, both plastic and nonplastic cases showed high host and virus population densities at low dilution rates. In both cases, low *w* led to lower nutrient availability and growth. The host lower growth associated with lower *w* selected for smaller cell radii, which showed reduced viral adsorption (Eq.(7)) [34]. This strategy allowed the host to reach high density levels, aided by the evolutionary strategy for the virus (increasing the duration of infection). The (classic) positive dependence between burst size and latent period observed for the nonplastic case, however, led to an increased viral pressure that in turn resulted in high viral densities at the expense of the host population; as a result, the host showed lower densities than in the plastic case. For the latter, the fact that plasticity breaks the classic *B*-*L* tradeoff means that the virus did not produce as much offspring per infection for low *w*, but the resulting increase in host density still enabled high viral densities; therefore, counterintuitively, due to viral plasticity, poor-growth conditions led to higher densities for both host and virus than in high-growth conditions [11]. The situation was maintained by a feedback loop in which increased host density led to lower nutrient availability and consequently host growth, decreasing viral performance and thus allowing for high host densities.

As the dilution rate increased, host size, growth, and adsorption all increased and allowed for shorter infections, which led to a lower host density. Although the burst size decreased for the nonplastic case and slightly increased for the plastic case, in both cases the viral population also decreased, which points to host availability (and not quality) driving this behavior. This decrease continued for the nonplastic case as *w* increased, as both *L* and *B* reached a lower value and thus the increase in dilution was not countered by an increase in viral production; increased dilution led to a decrease in viral density that in turn enabled an increase in host density for high dilution rates. For the plastic case, the latent period also reached a lower value but the increasing host growth enabled a higher burst size; as a result, the viral population increased with *w* for mid-to-low *N*_0_, whereas the lower *B* observed for high nutrient input led to a continuous decrease in viral availability. This lower viral density for high *w* and *N*_0_ does not result from the superinfection avoidance term, since similar behavior was observed for a version of the model with *α*_*V*_ = 0 (not shown). Host availability increased as dilution rate increased due to the improved growth conditions despite the improved viral performance, although higher *N*_0_ led to lower host densities as infection times shortened.

In summary, both plastic and nonplastic cases showed a convex dependence of host density on dilution, with a minimum of host availability for intermediate dilution rates resulting from the balance between bottom-up and top-down pressures. This minimum was thus not plasticity-driven, although accounting for plasticity led to higher host and viral densities.

### Coevolution vs single evolution

The convex trait curves comparing the outcome of coevolution with instances of single evolution obtained with this version of the model were in contrast with the concave curves obtained with the previous version [14]. The fact that the cell radius emerging from the coevolution with the virus transitioned from lower than to similar to the *r*_*ESS*_ obtained in the absence of the virus as *w* increased means that, under low-growth conditions, the outcome of host evolution was regulated by both nutrient availability and viral pressure; however, under high-growth conditions, top-down regulation played no significant evolutionary role. Nonetheless, high *N*_0_ values led to very large hosts in the absence of the virus, sizes that with the virus expose the host to viral infection but increase host maximum growth rate under decreased competition for resources (i.e. higher top-down and reduced bottom-up pressures); in this case, host evolution was thus both bottom-up and top-down regulated for any *w*. On the other hand, the convexity of the host density curve results from the eco-evolutionary dynamics with the virus, since the curve is instead concave in the absence of the virus (not shown).

For the virus, coevolution played an increasingly important role and shortened the emerging latent period as dilution rate and nutrient input increased. Given that the value in the absence of host coevolution, *L*_*nocoev*_, represents the latent period that “minimizes resources” for the virus, the departure from the 1:1 line indicates that other evolutionary strategies were prioritized by the virus under those environmental conditions. The dependence on *w* may stem from a shift from host availability dominating the viral evolutionary strategy when dilution is low, to host size and growth rate being the main influencing factor as *w* increases. This hypothesis is reinforced by the sensitivity of the curve to *N*_0_ for high dilution rates observed only for the plastic case. For high nutrient input levels, the latent periods emerging for plastic and nonplastic cases agreed, as host growth remained around its maximum across *w* and thus any plastic viral trait remained effectively fixed. Another relevant factor to consider is that for high nutrient levels, and for the nonplastic case, demographic oscillations are more marked and thus the most suitable viral strategy may not be the one that prioritizes the minimization of resources but one accounting for the changing environment, which has been observed in microbes such as phytoplankton [47].

### Evolutionary collapses

The modifications implemented in the current model enabled more dynamical stability than previous iterations [13, 14], as fewer initial conditions led to artificial (early) extinctions.

In addition to these early extinctions, we also observed evolutionary collapses. In these cases, the host population collapsed first, followed by the viral population. Also common to these cases, the initial random viral phenotype showed a high latent period (*L >* 1 *d*) and the sequence of dominant phenotypes was approaching a lower *L* at the moment of host extinction. Given the proximity of the host radius before extinction to the *r*_*ESS*_ reached without the virus, which we observed for many of these replicates, we speculate that extinction resulted from the host not being able to respond to changes in top-down regulation in a timely manner. The long infection times eased the evolutionary pressure on the host and, as a result, its evolving trait targeted the value expected under bottom-up regulation alone (i.e. the virus did not influence host evolution). As host and virus evolved, infection time decreased, increasing mortality on the host, which could not adapt fast enough to the environmental changes and went extinct. This hypothesis is reinforced by the fact that increasing the dilution rate led to lower *L* values dominating at the moment of extinction: higher dilution means that the abiotic component of the environment exerts a higher pressure on the host and, therefore, lysis time needs to be smaller for the mortality rate due to lysis to become important for host dynamics. In the plastic case, the situation may be exacerbated by high growth rates from the dominant host guaranteeing high burst sizes for the virus. The evolutionary collapse is in this case a combination of an “evolutionarily underperforming” host and an “ecologically overachieving” (plastic) virus.

All instances of coexistence showed a more parsimonious evolutionary path in which the initial viral phenotype showed a lower *L* regardless of the initial value of *r*, although a low initial *L* did not guarantee eventual coexistence as random initial extinctions were still possible.

## Conclusions

Our broader study of the coevolution of host size and viral latent period under chemostat conditions has confirmed that increasing dilution rates select for larger cell sizes and smaller latent periods, which is in agreement with the observation that better host quality and availability select for shorter infections [5, 48]. In our plastic description, these conditions also mean higher burst size, thus confirming biologically plausible strategies for both host and virus. For oligotrophic conditions, evolution leads to smaller host sizes that reduce viral adsorption, and a longer infection time allows the virus to compensate the handicapped host physiology by enabling the recovery of host density between infections. For eutrophic conditions, both host and virus can switch to more opportunistic strategies, with larger and faster-growing cells that allow the host population to survive even with the accompanying high adsorption rates, and with short and very productive lytic cycles that the virus can sustain thanks to the high host growth.

The information above helps understand the factors that regulate microbial communities. The flows present in a chemostat can roughly represent a volume of water in the ocean (with advection moving nutrients, microbes, and viruses in and out of the focal volume, and turbulence mixing the medium), or the directional flows present in the intestinal tract. The gut microbiome, for example, would be an example of high nutrient input and availability, and therefore our model predicts that the dominating radii in the bacterial community in the absence of the virus would be much larger than in the presence of phage. This could be relevant for predicting the fate of a viral infection affecting the microbiome. A similar situation would apply to nutrient-rich parts of the oceans, like cold or coastal waters, or zones of upwelling. In oligotrophic environments, on the other hand, the expectation would be for microbial hosts to show smaller sizes than in the absence of viruses (e.g. lab cultures), prediction reinforced by the increased competitive ability of small cells when taking up nutrients [49, 50]. Our eco-evolutionary model is especially well-suited to describe dynamic situations, like pulses affecting host growth temporarily [51], or when the distribution of resources translates into a spatial dependence for host growth rate, like in biofilms [26] or for soil bacteria-virus systems [52].

The evolutionary collapses we observed may provide useful information for the use of phages to eliminate bacterial infections (phage therapy) [25]. Our results indicate that infecting a bacterial population with a virus with a long latent period would result in a coevolutionary path in which hosts adapt their size mostly to bottom-up pressures due to the long lysis times, and ultimately will not be able to survive the combination of high burst size (due to the high host growth rate) and decreasing infection times. In our simulations, we did not observe replicates in which the viral population went extinct as a result of coevolution, although an alternative use of our model would be to predict how to use bacteria to eliminate viral infections.

One important limitation of our framework is that there are other potential targets for evolution in the system that may be subject to a higher pressure than host size and viral latent period. A situation that has been repeatedly studied in the past is the coevolutionary race that ensues when bacteria develop (total or partial) immunity against viral infection, and the virus tries to overcome such immunity [30, 53, 54]. One such examples is host avoidance of infection through a modification of LPS or other receptors, immunity that can be permanent or temporary depending on the viral capacity to, e.g. modify the tail fibers [42]. We are currently working on a version of the model that shifts the evolutionary focus towards changes in receptors by the host to avoid viral infection, and changes in host range by the virus that try to overcome such immunity and/or infect other host strains. We aim at understanding the role of plasticity in shaping this coevolutionary arms race.

The quantitative and qualitative differences observed here between the plastic and nonplastic versions of the model emphasize the need to consider plasticity in predictive theories for the eco-evolutionary dynamics of hosts and viruses. More empirical information is needed, however, to characterize viral plasticity in other important systems (e.g. different marine phytoplankton and virus species) and obtain the necessary expressions that unleash formalisms like the one presented here.

## Acknowledgments

The authors would like to thank the anonymous reviewers of this and previous publications for helpful suggestions. JAB was supported by a grant from the Simons Foundation (award #826106); MC and MRH were supported to varying extents by the Marine Alliance for Science and Technology for Scotland (MASTS) pooling scheme, Scottish Funding Council grant reference HR09011.

## Data availability

No relevant data other than the equations above were used in this research. Code used to implement such equations is available without restriction under request to the corresponding author.

## Supplementary Information

### Host-use minimization strategy when only virus evolves

It is possible to calculate analytically the optimal latent period of a virus that evolves trying to minimize the requirement of a (nonevolving) host. In other words, the latent period that minimizes the density of hosts needed for the survival of the virus phenotype. This would be equivalent to finding the virus with the smallest “*R*^*^”, using the jargon of resource competition theory [40].

Assuming that the dynamics of the system follow Eqs.(1)-(4), setting the equations to zero provides the condition for the stationary state. For example, from setting the r.h.s. of Eqs.(1) and (3) to zero, we obtain the expression:

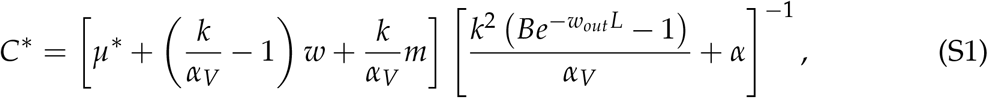

where the asterisk represents “value at the stationary state”. Because the host does not evolve, the value of the latent period that minimizes the expression given by Eq.(S1) needs to fulfill:

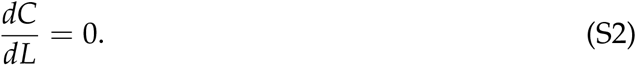

Given the expression for *C*^*^ above, such a condition is fulfilled if the derivative of the denominator in Eq.(S1) is zero, which ultimately leads to the condition:

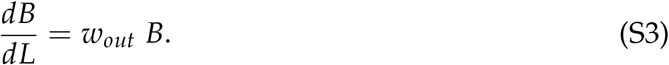

Since the burst size is linked to the latent period through the tradeoff function Eq.(9), the condition above translates into:

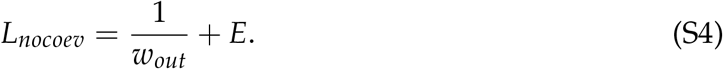

Thus, Eq.(S4) describes the evolutionarily stable strategy (ESS) for a virus for which evolution aims to minimize the host density needed for survival, in the presence of a host that does not evolve. This expression is identical to the ones obtained in the past for other versions of the delay model when calculating the optimal latent period of the virus [11, 29, 44].

## Supplementary Figures

**Figure S1:**
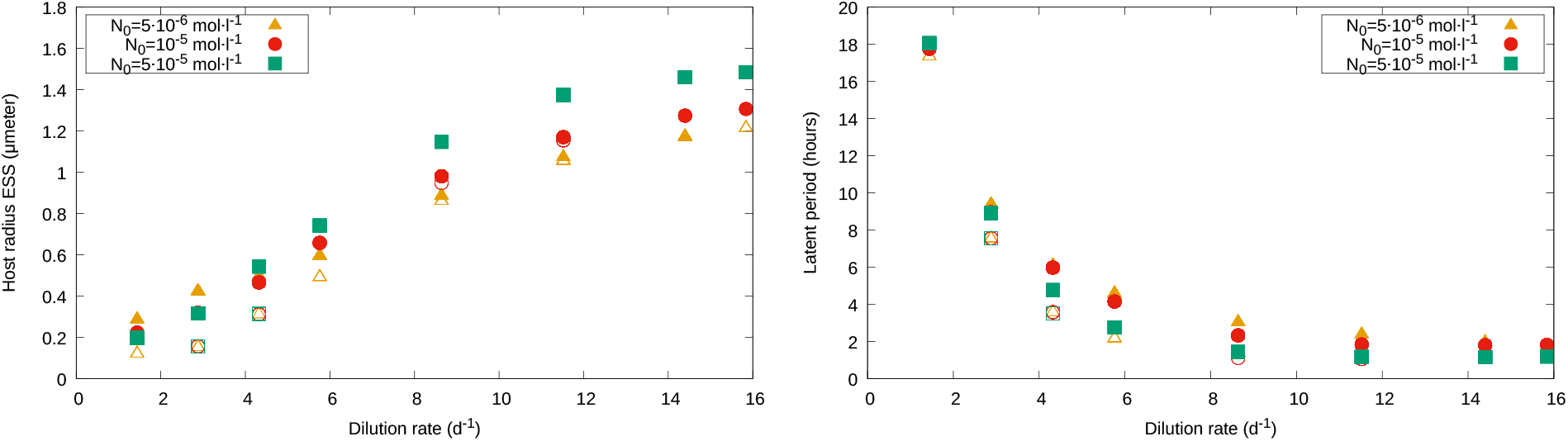
Dependence of the evolving traits on the dilution rate, *w*. The emergent *r*_*ESS*_ value (left) increases with *w* and input concentration, *N*_0_, whereas the latent period decreases (right).

**Figure S2:**
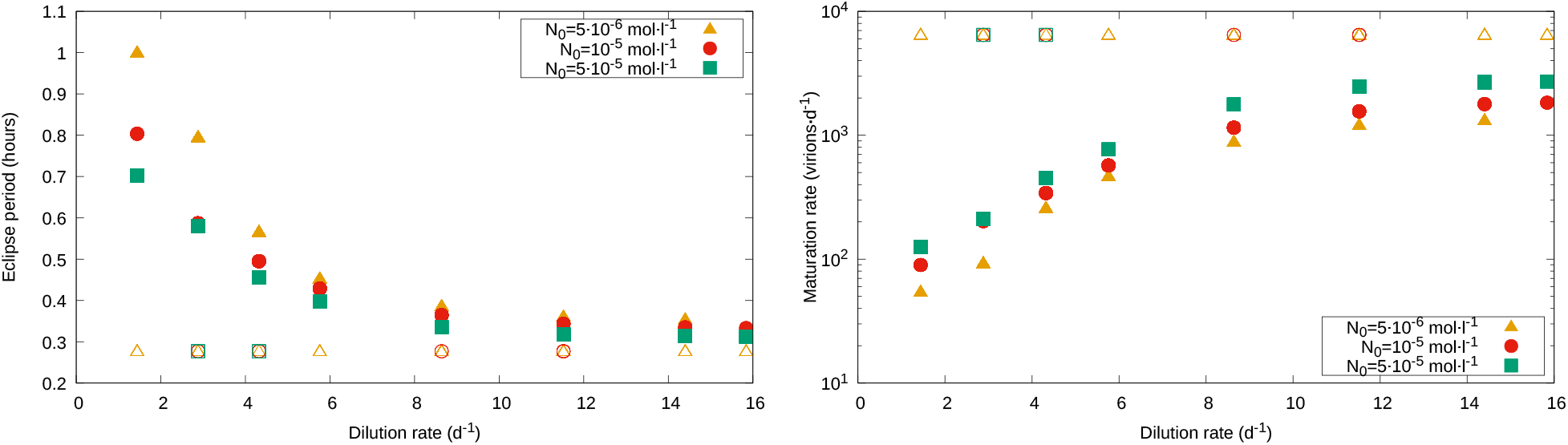
Eclipse period (left) and maturation rate (right) as a function of the dilution rate. Both are constant for the nonplastic case, but depend on growth rate in a negative and positive way, respectively, in the plastic case.

**Figure S3:**
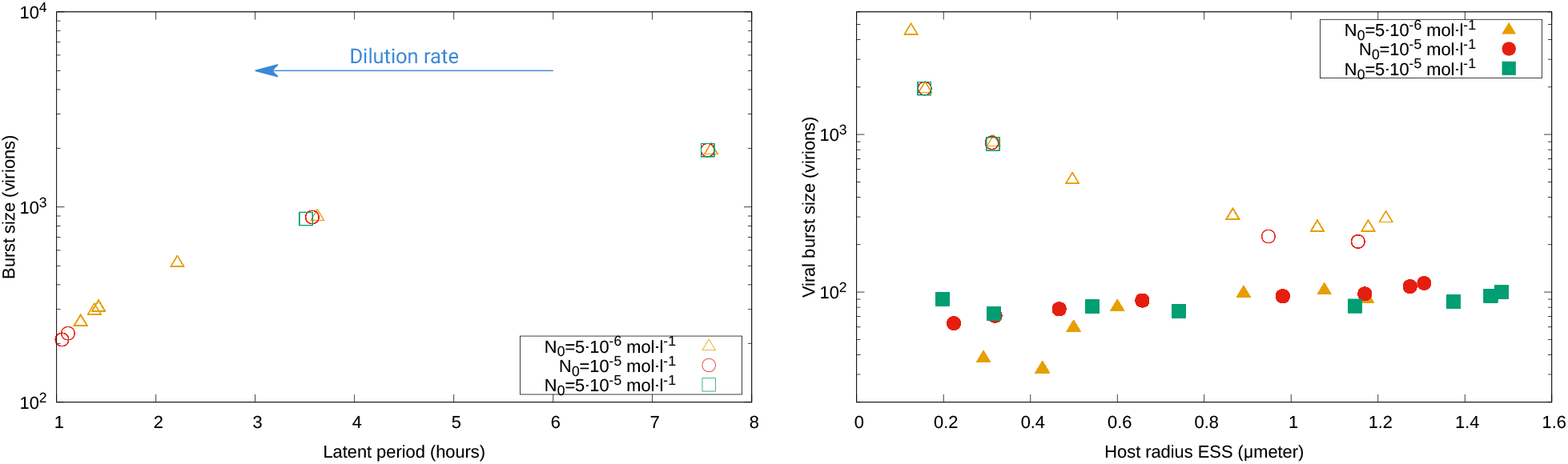
Left: Burst size as a function of the emerging latent period for the nonplastic case, which shows a positive correlation as expected from using Eq.(9) with fixed *E* and *M*, and the nonplastic *L*_*ESS*_ from Fig. 3, left panel. Right: Average across replicates of emerging burst size as a function of host radius; the plastic case shows a mild positive correlation, whereas the nonplastic version shows a marked negative correlation.

**Figure S4:**
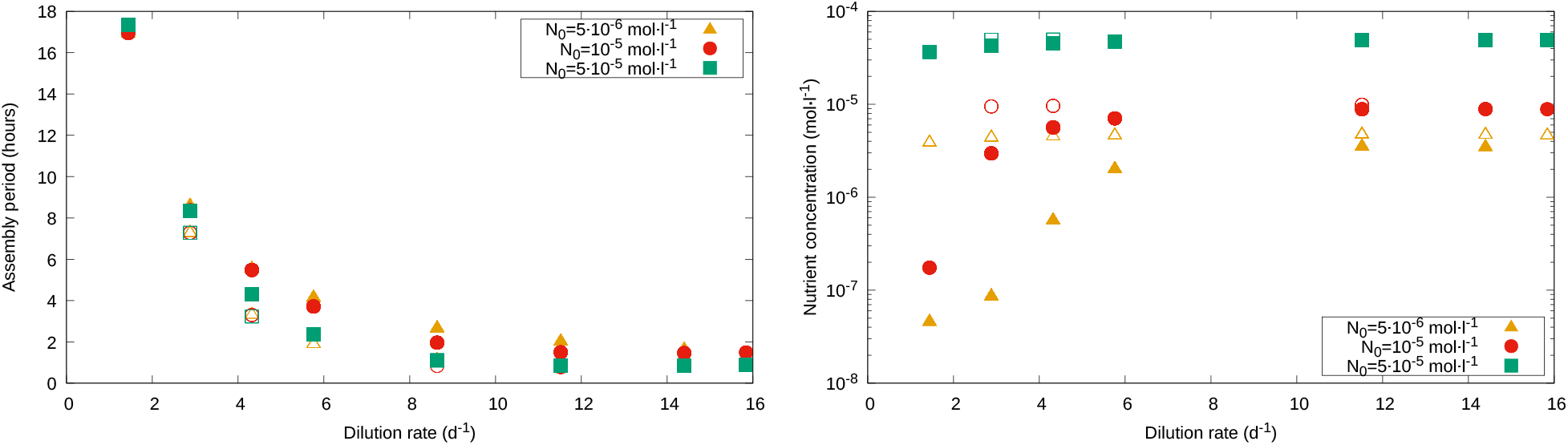
Left: Assembly period (time between end of eclipse period and lysis, *L* − *E*) as a function of dilution rate. Right: Average across replicates of nutrient concentration as a function of *w*; the plastic case shows lower *N* for low dilution rates, but both converge to the input concentration *N*_0_ as *w* increases.

**Figure S5:**
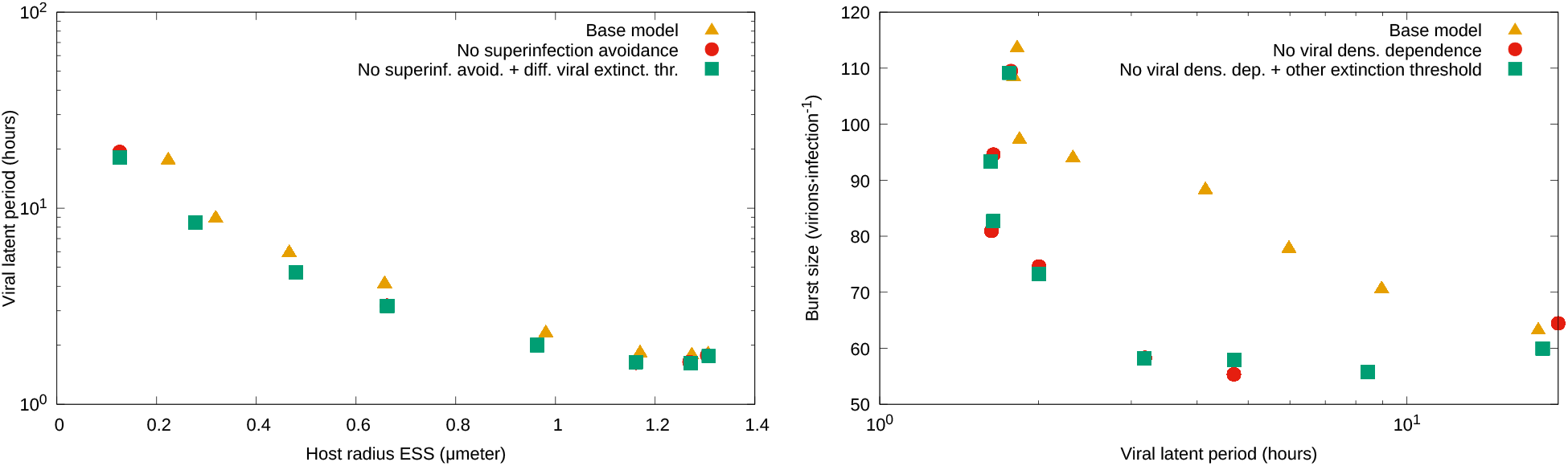
Emerging traits for other versions of the plastic model (see main text). Left: Host radius as a function of viral latent period, for which the base model (i.e. main model described in Methods) shows results that are barely distinguishable from the case with no superinfection avoidance (i.e. with *α*_*V*_ = 0) and from the case with a threshold viral for extinction that depends only on free virus availability. Right: Burst size as a function of the latent period, which shows a qualitatively similar decreasing trend for all three versions, but the base model produces longer infections with larger burst sizes.

**Figure S6:**
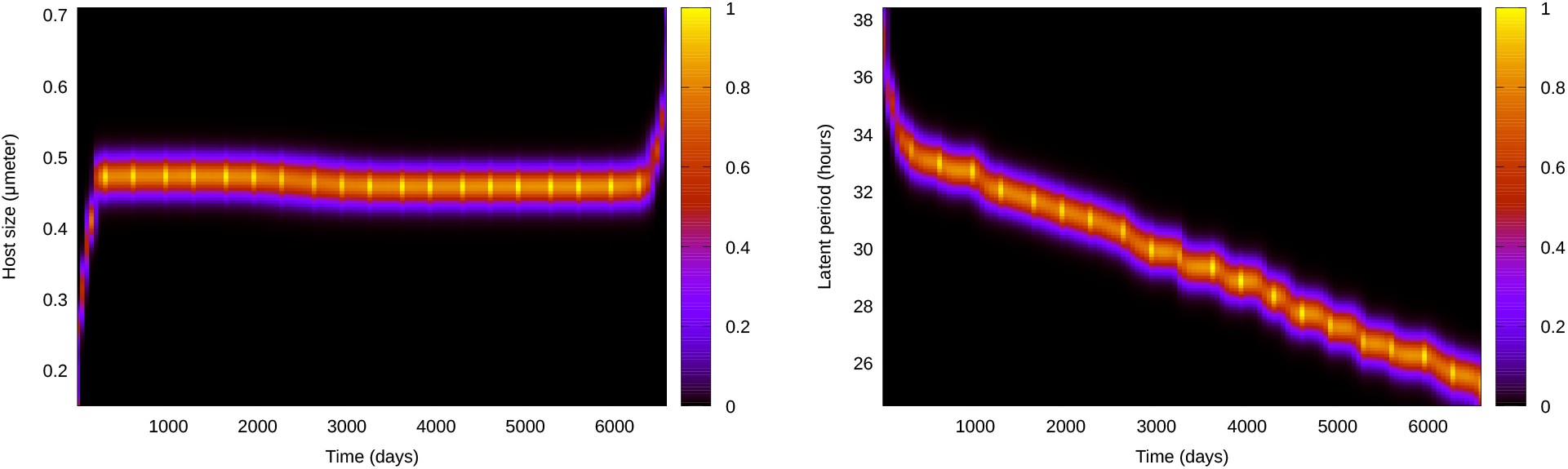
Evolutionary paths to system collapse for the plastic case with *w* = 10, *N*_0_ = 5·10^−5^. Left: Host radius, *r*. Right: Viral latent period, *L*.

**Figure S7:**
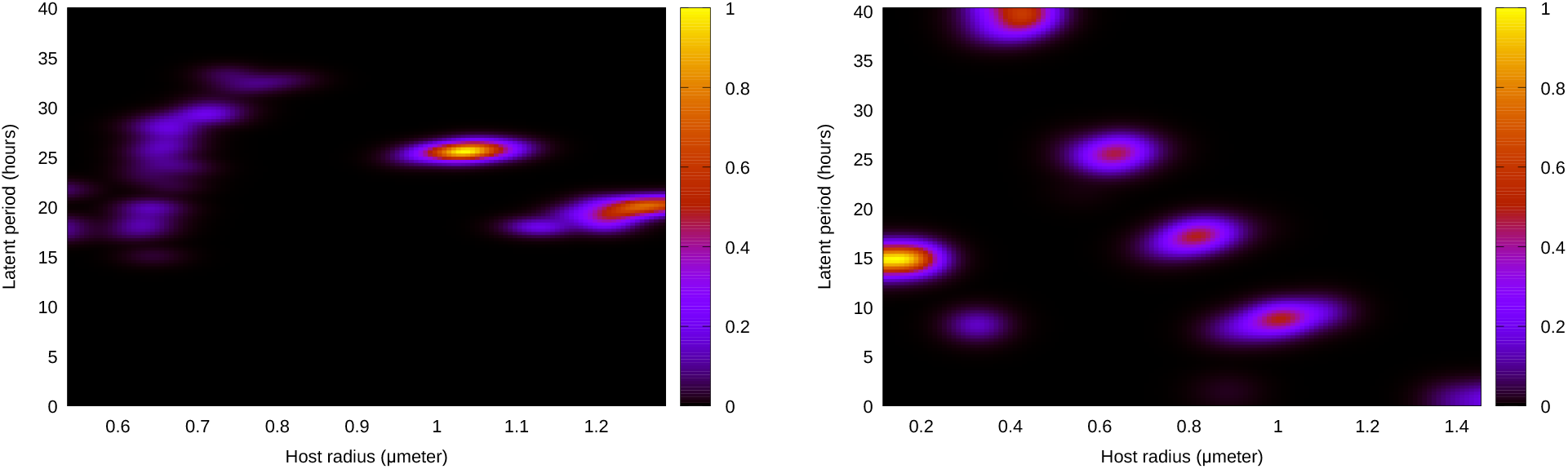
Representative examples of (normalized) density plots for the host radius and viral latent period observed immediately before evolutionary collapse of the system occurred for the plastic (*N*_0_ = 5 · 10^−5^ *mol · l*^−1^, left) and the nonplastic (*N*_0_ = 10^−5^ *mol · l*^−1^, right) case. See Fig. S6 for examples of evolutionary path to extinction for the plastic case. As in Fig. 2, dilution rate increases from left to right.

